# Detection of cell-type-specific risk-CpG sites in epigenome-wide association studies

**DOI:** 10.1101/415109

**Authors:** Xiangyu Luo, Can Yang, Yingying Wei

## Abstract

In epigenome-wide association studies, the measured signals for each sample are a mixture of methylation profiles from different cell types. The current approaches to the association detection only claim whether a cytosine-phosphate-guanine (CpG) site is associated with the phenotype or not, but they cannot determine the cell type in which the risk-CpG site is affected by the phenotype. Here, we propose a solid statistical method, HIgh REsolution (HIRE), which not only substantially improves the power of association detection at the aggregated level as compared to the existing methods but also enables the detection of risk-CpG sites for individual cell types.

---

Epigenome-wide association studies (EWAS) aim to identify cytosine-phosphate-guanine (CpG) sites associated with phenotypes of interest, for example, disease status [1, 2, 3], smoking history [4, 5], body mass index [6], and age [7, 8]. However, as samples in EWAS are measured at the bulk level rather than at the single-cell level, the obtained methylome for each sample shows the signals aggregated from distinct cell types [3, 9, 10], leading to two main challenges for analyzing EWAS data. On the one hand, the cell type compositions differ between samples and can be associated with phenotypes [3, 10]. Both binary phenotypes, such as the diseased or normal status [3], and continuous phenotypes, for example, age [10], have been found to affect the cell type compositions. As a result, ignoring the cellular heterogeneity in EWAS can lead to a large number of spurious associations [10, 11, 12, 13]. On the other hand, the phenotype may change the methylation level of a CpG site in some but not all of the cell types. Identifying the exact cell types that carry the risk-CpG sites can deepen our understandings of disease mechanisms. Nevertheless, such identification is challenging because we can only observe the aggregated-level signals.

To the best of our knowledge, no existing statistical method for EWAS can detect cell-type-specific associations despite the active research on accounting for cell-type heterogeneity. The existing approaches can be categorized into two schools [14]: “reference-based” and “reference-free” methods. The reference-based methods [9, 15] require the reference methylation profiles for each cell type to be known a priori, and they regress the aggregated methylation levels observed from each sample on the same set of references to learn the sample’s cellular compositions. However, as samples have different attributes, such as age and gender, the methylation levels of a given cell type can vary with samples. Therefore, it is problematic to assume that all of the samples have the same set of reference profiles [10, 14]. Furthermore, high-quality references are difficult to obtain for most EWAS due to the existence of unknown cell types, the high cost of cell sorting, and confounding effects [14]. Consequently, a large amount of recent EWAS literature was devoted to finding risk-CpG sites without the need of the reference methylation profiles.

The reference-free methods in general can be further divided into two classes according to whether they estimate the cell-type mixing proportions directly or not. The direct-decomposition-based procedures consist of the following two stages: in the first stage, they simultaneously estimate the cellular compositions for each sample and the cell-type-specific reference methylomes via quadratic programming [16]; and in the second stage, the direct-decomposition-based methods treat the estimated cell-type proportions as covariates with additive effects in the linear models to conduct association tests. However, when estimating cellular compositions in the first stage, the direct-decomposition-based methods also do not consider samples’ phenotype information, thus suffering from the same problem of biasing the cellular composition estimates as the reference-based approaches [9]. Moreover, similar to tumor purity [17], we argue that the estimated cellular composition has a *multiplicative* rather than *additive* effect on the observed methylation level (Methods). The second class of methods, exemplified by SVA [18], RefFreeEWAS [19], and ReFACTor [13], do not carry out cell-type decompositions. They resort to singular value decomposition, which includes the principal component (PC) analysis, to construct surrogates for the underlying cell-type composition. EWASher, a linear mixed model, also belongs to this class as it is “equivalent to using PCs as fixed effect covariates” [11]. However, the use of PCs as the covariates in the regression undergoes the same issue of additive effects as the direct-decomposition-based methods. Therefore, the existing reference-free methods have low power in detecting risk-CpG sites [12].

Although all of the existing methods aim to address the cellular heterogeneity problem in EWAS and claim whether a CpG site is associated with phenotypes at the *aggregated level*, none of them can identify the risk-CpG sites for each *individual cell type*, thus missing the opportunity to obtain finer-grained results in EWAS.

Here, we propose a new method HIRE to identify the association in EWAS at an un-precedented HIgh REsolution: detecting whether a CpG site has any associations with the phenotypes in *each cell type* (Methods). The keys to HIRE’s success are twofold. First, HIRE links the underlying cell-type-specific methylation profiles for each sample to the sample’s phenotypes, thus avoiding the bias in estimating the cellular composition by the reference-based and direct-decomposition-based methods. Second, HIRE correctly characterizes the cellular compositions as the multiplicative effects, whereas the cell proportions are inappropriately treated as additive effects by the existing methods (Methods). HIRE is applicable to EWAS with binary phenotypes, continuous phenotypes, or both. By helping researchers understand in which cell types the CpG sites are affected by a disease, HIRE can utlimately facilitate the development of epigenetic therapies by targeting the specifically affected cell types.

## Results

### Method overview

HIRE is a hierarchical model that closely follows the data generation process. Its elaborate modeling depicts how phenotypes affect the methylation levels of each sample. Here we briefly introduce the method. The technical details are provided in the Methods section and the Supplementary Note.

To begin with, let us review the cornerstone in most EWAS approaches. These methods model the observed methylation levels of the *m* CpG sites for sample *i*, **O**_*i*_ = (*O*_1*i*_, *O*_2*i*_*,…, O*_*mi*_)^*T*^, as the weighted average of the methylation profiles of *K* cell types, **u**_*i*_ = (**u**_*i*1_, **u**_*i*2_*,…,* **u**_*iK*_). The weights are the cellular compositions **p**_*i*_ = (*p*_1*i*_, *p*_2*i*_*,…, p*_*Ki*_)^*T*^ of sample *i* (see the top panel of Fig. 1a). However, no matter the reference is known a priori or not, the existing methods assume that the cell-type-specific methylation profiles **u**_*i*_s stay the same for all of the samples: **u**_*i*_ = **M**, for *i* = 1, *…, n*. Unfortunately, because methylation levels can actually change with covariates such as age and disease status, ignoring the covariates effects and enforcing static reference methylomes can bias the estimation of **p**_*i*_ and subsequently affect all the downstream analyses [14]. More importantly, assuming that cell-type-specific methylation profiles are the same for each sample forbids the detection of cell-type-specific risk-CpG sites.

**Figure 1:**
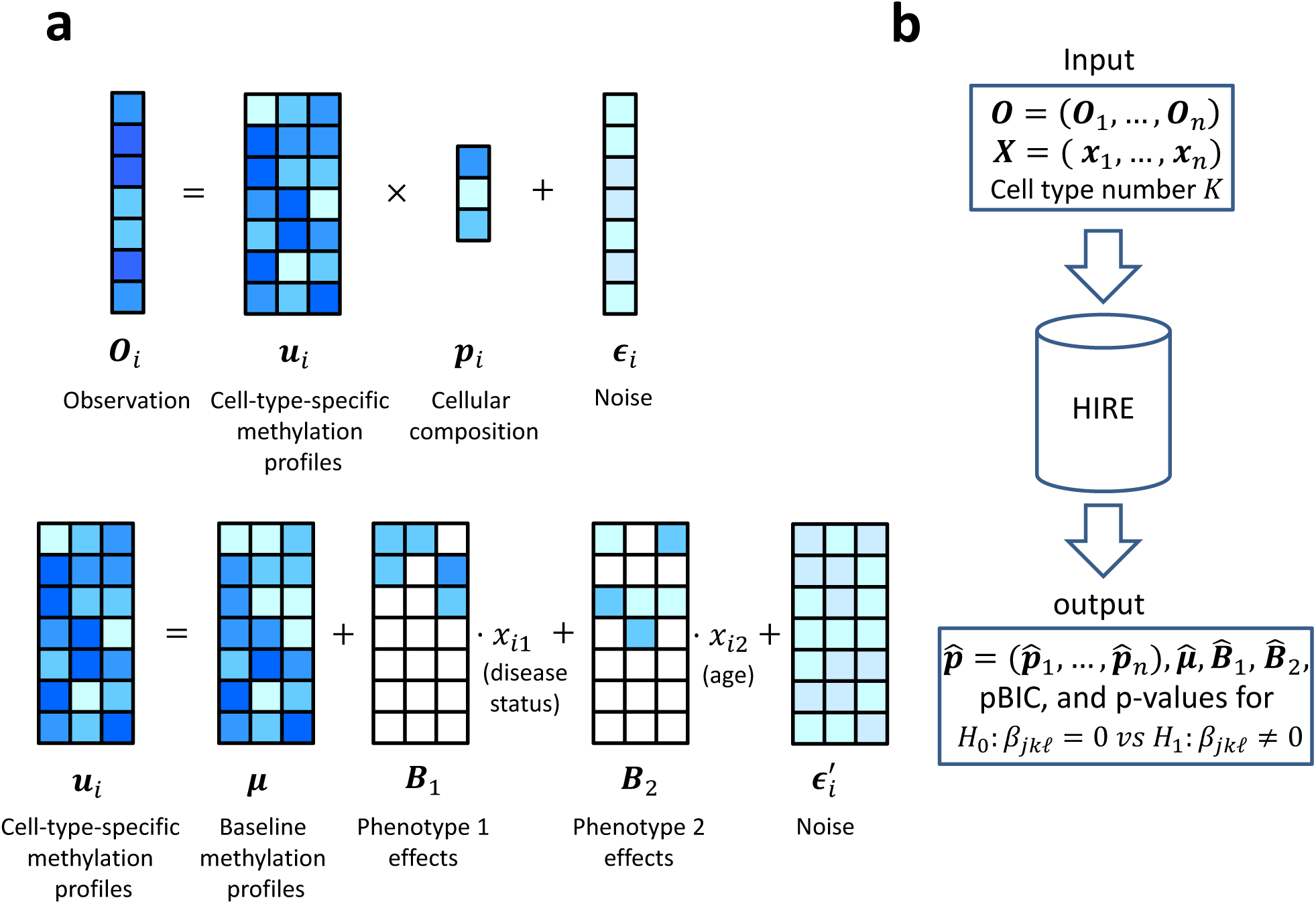
A simple cartoon illustration of the HIRE model, where there are three cell types (*K* = 3) and two phenotypes, disease status and age (*q* = 2). (**a**) The data generation procedure for the observed methylation vector **O**_*i*_ for sample *i* (*i* = 1*,…, n*). In the top panel, **O**_*i*_ is the convolution of cell-type-specific methylation profiles **u**_*i*_ with cellular compositions **p**_*i*_. Both **u**_*i*_ and **p**_*i*_ depend on the attributes of sample *i*. The bottom panel describes how sample *i*’s phenotypes affect **u**_*i*_ through two phenotype-effect matrices **B**_1_ and **B**_2_. In **B**_1_ and **B**_2_, the white square represents zero, which means that the phenotype has no influence on the corresponding methylation level in **u**_*i*_. (**b**) The inputs and outputs of HIRE. We input the observed methylation matrix **O**, the phenotype data matrix **X**, and a predetermined cell type number *K* into HIRE, and HIRE outputs the estimates for cellular compositions 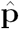, baseline methylation profiles 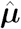, phenotype effects 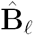, and the penalized BIC value. In addition, HIRE tests whether there is any association between CpG site *j* and phenotype *£* in cell type *k*—*H*_0_ : *β*_*jk*_ = 0 vs *H*_1_ : *β*_*jk*_ *≠* 0—and provides the p-values.

For association detection at the aggregated level, after estimating **p**_*i*_ using the deconvolution-based approach or its surrogates from PC-based methods, the existing methods examine a linear model where the phenotypes **x**_*i*_ = (*x*_*i*1_*,…, x*_*i*_ *,…, x*_*iq*_)^*T*^ and the cellular proportions **p**_*i*_ have additive effects on the methylation level **O**_*i*_:

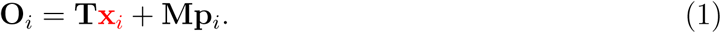

Subsequently, a CpG-site *j* is associated with phenotype *𝓁* if we reject the null hypothesis that the covariate coefficient *T*_*j*_ equals zero.

In contrast, HIRE further models the impact of each phenotype on each cell type as shown in the bottom panel of Fig. 1a. In cell type *k*, sample *i*’s cell-type-specific methylation profile, **u**_*ik*_, is the summation of the corresponding baseline cell-type-specific methylation levels, ***µ***_*k*_, and the phenotype effects **B**_*k*_ *x*_*i*_ on sample *i* from all the *l* = 1, *…, q* phenotypes: 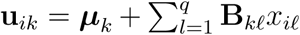, where *x*_*i*_ is the phenotype *𝓁* of sample *i* and **B**_*k*_ = (*β*_1*k*_ *,…, β*_*mk)*_^*T*^ — the *k*th column of **B** —reflects the association of phenotype *𝓁* with each of the *m* CpG sites in cell type *k*. Consequently, by collecting the baseline cell-type-specific methylation profiles to ***µ*** = (***µ***_1_*,…,* ***µ***_*k)*_ and denoting the *m* by *K* phenotype coefficient matrix (*β*_*jk*_ : 1 *≤ j ≤ m,* 1 *≤ k ≤ K*) by **B**, now we have:

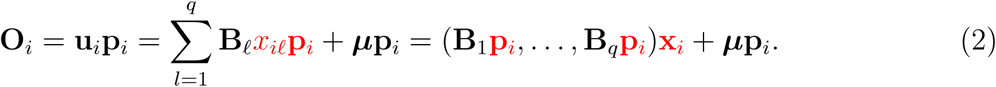

Comparing the red parts in Equation 1 and 2, we can see that via the two-layer hierarchical model HIRE correctly captures the multiplicative effects of the cellular compositions on the phenotype effects (see also Methods and the Supplementary Note). As a result, HIRE achieves an increased statistical power for the association detection at the aggregated level and enables the fine-scale resolutions that had previously been infeasible.

Fig. 1b summarizes the inputs and outputs of HIRE. Given the methylation measurements at the aggregated level of *n* samples, HIRE is able to estimate all the parameters of interests —**p**_*i*_ (*i* = 1*,…, n*), ***µ***, and **B** (*𝓁* = 1*,…, q*). Subsequently, HIRE determines whether there is any association between CpG site *j* and phenotype *𝓁* in each individual cell type by testing the hypotheses *H*_0_ : *β*_*jk*_ = 0 versus *H*_1_ : *β*_*jk*_ *≠* 0. When the null hypothesis *H*_0_ : *β*_*jk*_ = 0 is rejected, HIRE calls CpG site *j* as a risk-CpG site for phenotype *𝓁* in cell type *k*. The detection of cell-type-specific risk-CpG sites is not available by all the existing state-of-the-art methods. Moreover, HIRE allows the users to pre-specify the number of cell types *K*. When *K* is unknown, HIRE selects the cell type number according to the penalized Bayesian information criterion (pBIC) [20] (Supplementary Note).

### Simulation

As the definition of the gold standards for real data is debatable [21, 22], we designed extensive simulation studies to evaluate the performance of HIRE and compared it with commonly used methods—unadjusted analysis, SVA, RefFreeEWAS, EWASHer, and ReFACTor (Methods). We generated datasets where the observed methylation was a mixture of several cell types and each sample was accompanied with a diseased or normal status and a continuous age attribute. We deliberately designed some cell types with similar baseline methylation profiles to mimic cell types from the same cell lineage. We set the sample size *n* to 180, 300, and 600 and let the underlying cell type number *K* be 3, 5, and 7. For each pair of (*n, K*), we investigated two scenarios: (1) all of the phenotype effects *β*_*jk*_ s are zero—the “*true null* ” case—to compare each method’s ability to control false positives; and (2) a small portion of *β*_*jk*_ s are non-zero—the “*true alternative*” case—to study the power of each method for detecting risk-CpG sites. Under the “true alternative,” both the binary and the continuous phenotypes were assumed to have cell-type-specific risk-CpG sites and to affect the cell-type proportions among the samples [10]. We further simulated phenotype effects with different directions and magnitudes.

Under the “true null,” HIRE, EWASHer, and ReFACTor control the FPR very well, none of which are greater than 0.05% (Table 1 and Supplementary Figs. 1-9). In comparison, RefFreeEWAS often has FPRs larger than 0.1%, performing not as well as HIRE, and the unadjusted analysis and SVA further suffer from the dramatic inflation of false positives. For the “true alternative” settings, given that the FPRs are well-controlled, with FPRs below 0.05%, HIRE achieves the highest TPR among all of the methods in every simulation setting (see also Figure 2a and Supplementary Figs. 10-17). As expected, as the sample size increases, HIRE’s power increases. For example, when there are five cell types in the data, HIRE can identify 89.6% of the risk-CpG sites with 300 samples, and HIRE is able to detect almost all of the risk-CpG sites when the sample size reaches 600, which is a typical sample size for an EWAS. Although EWASHer and ReFACTor have low FPRs, they miss a large proportion of risk-CpG sites. EWASHer’s maximum TPR is only 35.33%, and ReFACTor’s maximum TPR is slightly more than 60%. However, in those cases, HIRE’s power is greater than 95%. Consistent with the “true null” scenario, in the “true alternative,” RefFreeEWAS has inflated FPRs compared to HIRE, and the unadjusted analysis and SVA always have explosive false positives. Therefore, HIRE substantially improves the power of association detection at the aggregated level compared with existing methods.

**Table 1:**
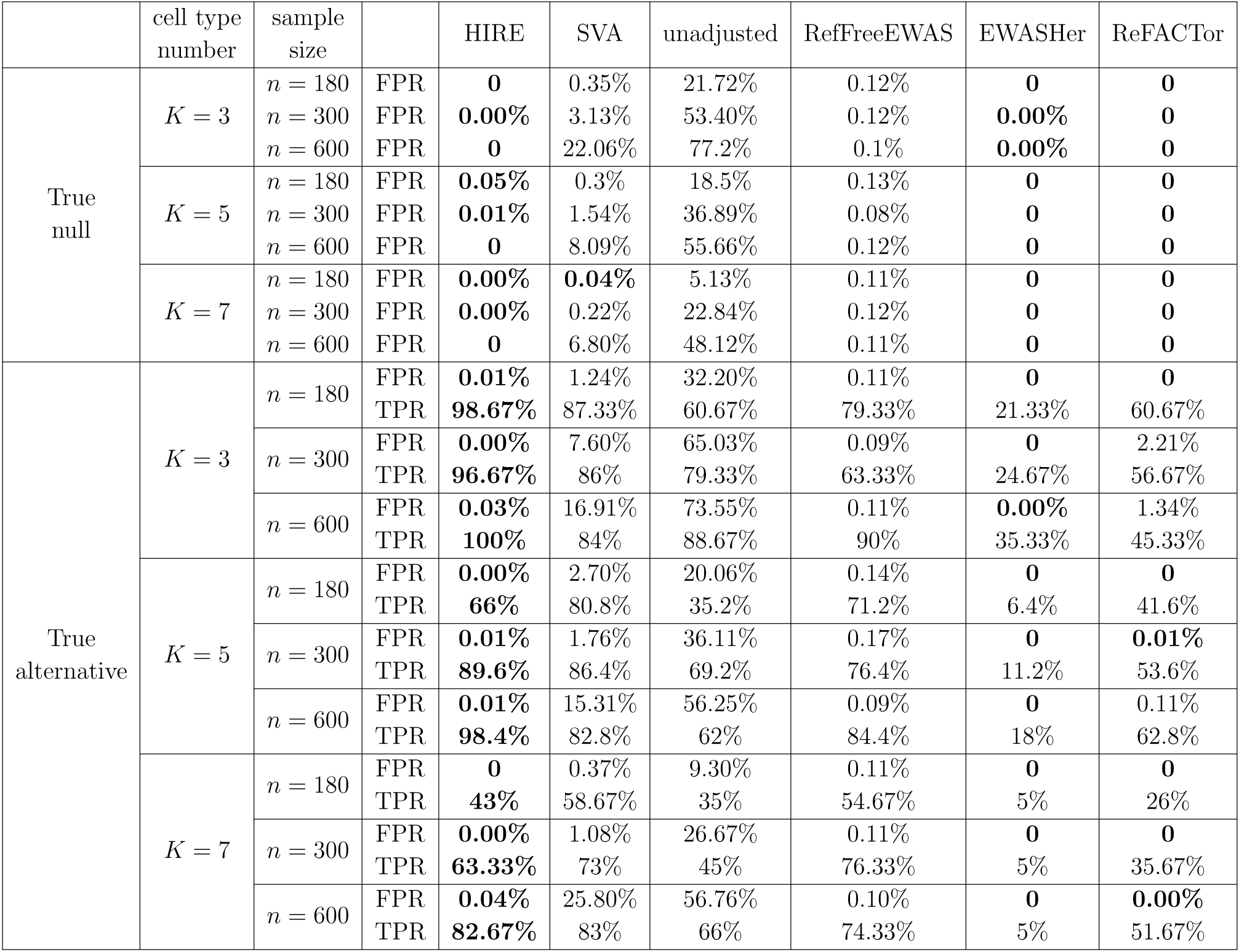
The performance of HIRE, SVA, unadjusted analysis, RefFreeEWAS, EWASHer and ReFACTor in simulation studies in detecting risk-CpG sites at the aggregated level. For the “true null” cases where no CpG site is at risk, the average of the false positive rates (FPRs) based on five replicates is reported. For the “true alternative” cases, the averages of the FPRs and the true positive rates (TPRs) based on five replicates are reported. The number of CpG sites at risk is 30, 50, and 60 for the cell type number *K* = 3, 5, and 7, respectively. HIRE calls a CpG site as significant at the aggregated level if it is at risk in at least one cell type. For all of the methods, we used Bonferroni correction to control the family-wise error rate (FWER) to be less than *α* = 0.01. For HIRE, since it can provide p-values of CpG sites for all cell types and phenotypes, the p-value threshold to call “significant” is *α/*(*mKq*), where *m* = 10, 000 is the number of CpG sites and *q* = 2 is the phenotype number. For the other five methods, the p-value threshold is set to *α/m*. FPRs less than or equal to 0.05% are shown in bold. In the “true alternative” settings, the maximum TPRs with FPRs controlling at 0.05% are highlighted. Notice that “0” represents exact zero, and “0.00%” indicates a very small positive number that is rounded to zero using four decimal places.

**Figure 2:**
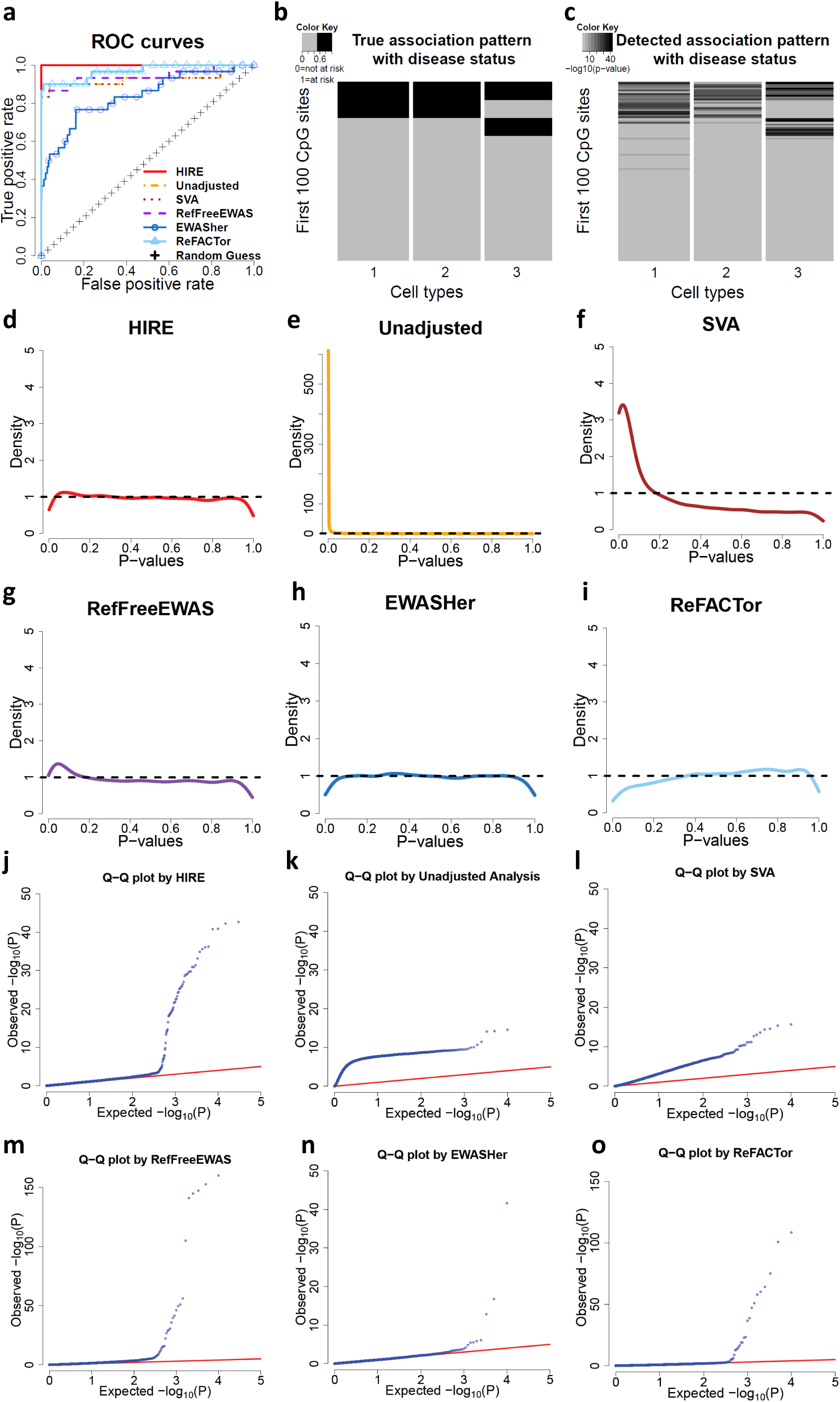
The association detection performance by HIRE and the commonly used methods in the “true alternative” setting with *K* = 3 and *n* = 180. (**a**) The ROC curves of HIRE and the commonly used methods. HIRE has the largest area under the curve among all of the methods. (**b**) The true cell-type-specific association pattern with disease status for the 10,000 simulated CpG sites, where the columns correspond to cell types and the rows represent the CpG sites. The dark cells correspond to risk-CpG sites, whereas the grey cells are CpG sites not associated with the disease status. (**c**) The detected cell-type-specific association pattern with disease status by HIRE. The darkness represents *-* log_10_(*p - value*) (**d-i**) The p-value density plots for association with disease status in the simulation dataset for (**d**) HIRE, (**e**) unadjusted analysis, (**f**) SVA, (**g**) RefFreeEWAS, (**h**) EWASHer, and (**i**) ReFACTor. (**j-o**) The Q-Q plots for association with disease status for (**j**) HIRE, (**k**) unadjusted analysis, (**l**) SVA, (**m**) RefFreeEWAS, (**n**) EWASHer, and (**o**) ReFACTor.

In the multiple hypothesis testing, the p-values from the truly null features should follow a uniform distribution on (0, 1), whereas those for the truly alternative features concentrate near zero [23]. Both the histograms (Figs 2d-i) and Q-Q plots (Figs. 2j-o) show that the p-value distribution of HIRE is the best fit to the underlying truth—there are only a small proportion of signals, followed by RefFreeEWAS and ReFACTor. EWASHer easily overcorrects signals with its p-value density having a dip near zero (Figure 2h), thus failing to detect the true associations. In contrast, the unadjusted analysis and SVA generate very small p-values clustered near zero, resulting in inflated type I errors.

In addition to the traditional association detection at the aggregated level, HIRE is able to identify the association for each CpG site with the phenotypes under each cell type. Table 2 shows the FPR and TPR of HIRE in each cell type for different simulation settings. Such fine analysis is not available from the other methods. Consistent with the association detection at the aggregated level, HIRE always controls the FPR well. When *K* = 3 and *n* = 180, HIRE accurately detects the risk-CpG sites associated with the disease status with the TPR more than 83% and the FPR less than or equal to 0.01% in all of the three cell types. Similarly, most of the CpG sites affected by age are also correctly identified in each cell type. HIRE’s learned cell-type-specific association patterns closely matches the underlying true associations (see Figs. 2b-c and Supplementary Figs. 18-26). Once again, HIRE’s power decreases with the number of cell types and increases with the sample size. When the samples consist of 7 cell types and the proportion of the least abundant cell type is as low as 4.2%, given a typical current EWAS with around 600 samples, HIRE can detect most cell-type-specific risk-CpG sites reasonably well. Moreover, HIRE’s estimates for the cellular compositions and phenotype effects have little bias (Supplementary Figs. 27-62). Therefore, HIRE can provide accurate estimates and is powerful in detecting cell-type-specific risk-CpG sites.

**Table 2:** The performance of HIRE in detecting cell-type-specific risk-CpG sites in the “true alternative” cases. The results are based on five replicates for each setting. A CpG site is claimed to be “significant” in a given cell type if its p-value is less than *α/*(*mKq*).

| Phenotype | cell type number | sample size |  | cell types |  |  |  |  |  |  |
| --- | --- | --- | --- | --- | --- | --- | --- | --- | --- | --- |
|  |  |  |  | 1 | 2 | 3 | 4 | 5 | 6 | 7 |
| Disease status | $K = 3$ | $n = 180$ | FPR | 0.01% | 0.00% | 0.01% | | | | |
|  |  |  | TPR | 83% | 85% | 92% |  |  |  |  |
| | | $n = 300$ | FPR | 0.02% | 0.02% | 0.04% | | | | |
|  |  |  | TPR | 74% | 85% | 95% |  |  |  |  |
| | | $n = 600$ | FPR | 0.03% | 0.03% | 0.05% | | | | |
|  |  |  | TPR | 99% | 98% | 100% |  |  |  |  |
| | $K = 5$ | $n = 180$ | FPR | 0.01% | 0 | 0.00% | 0.01% | 0.02% | | |
|  |  |  | TPR | 35% | 46% | 44% | 39% | 75% |  |  |
| | | $n = 300$ | FPR | 0.02% | 0.02% | 0.02% | 0.06% | 0.10% | | |
|  |  |  | TPR | 66% | 73% | 67% | 43% | 43% |  |  |
| | | $n = 600$ | FPR | 0.02% | 0.02% | 0.01% | 0.10% | 0.12% | | |
|  |  |  | TPR | 81% | 77% | 92% | 52% | 56% |  |  |
| | $K = 7$ | $n = 180$ | FPR | 0 | 0 | 0.01% | 0 | 0.00% | 0 | 0.00% |
|  |  |  | TPR | 13% | 28% | 32% | 20% | 21% | 15% | 69% |
| | | $n = 300$ | FPR | 0.01% | 0.01% | 0.01% | 0.00% | 0.01% | 0.01% | 0.02% |
|  |  |  | TPR | 20% | 48% | 60% | 52% | 40% | 23% | 78% |
| | | $n = 600$ | FPR | 0.02% | 0.02% | 0.01% | 0.02% | 0.01% | 0.01% | 0.07% |
|  |  |  | TPR | 37% | 79% | 90% | 52% | 71% | 66% | 98% |
| Age | $K = 3$ | $n = 180$ | FPR | 0.01% | 0.01% | 0.06% | | | | |
|  |  |  | TPR | 68% | 76% | 96% |  |  |  |  |
| | | $n = 300$ | FPR | 0.05% | 0.03% | 0.08% | | | | |
|  |  |  | TPR | 95% | 95% | 90% |  |  |  |  |
| | | $n = 600$ | FPR | 0.06% | 0.06% | 0.08% | | | | |
|  |  |  | TPR | 94% | 99% | 95% |  |  |  |  |
| | $K = 5$ | $n = 180$ | FPR | 0.05% | 0.05% | 0.01% | 0.04% | 0.06% | | |
|  |  |  | TPR | 67% | 61% | 82% | 69% | 97% |  |  |
| | | $n = 300$ | FPR | 0.09% | 0.03% | 0.04% | 0.04% | 0.08% | | |
|  |  |  | TPR | 78% | 85% | 97% | 85% | 91% |  |  |
| | | $n = 600$ | FPR | 0.07% | 0.06% | 0.07% | 0.08% | 0.08% | | |
|  |  |  | TPR | 88% | 84% | 94% | 83% | 94% |  |  |
| | $K = 7$ | $n = 180$ | FPR | 0.02% | 0.01% | 0.01% | 0 | 0.02% | 0.01% | 0 |
|  |  |  | TPR | 39% | 62% | 58% | 68% | 38% | 54% | 85% |
| | | $n = 300$ | FPR | 0.08% | 0.01% | 0.04% | 0.01% | 0.03% | 0.03% | 0.02% |
|  |  |  | TPR | 46% | 62% | 79% | 84% | 79% | 73% | 93% |
| | | $n = 600$ | FPR | 0.09% | 0.08% | 0.04% | 0.10% | 0.03% | 0.05% | 0.07% |
|  |  |  | TPR | 52% | 77% | 85% | 84% | 77% | 79% | 84% |

To further evaluate HIRE’s performance on experimentally mixed samples, we conducted another semi-simulated dataset, which includes six samples mixed with six purified cell types at pre-determined proportions [24]. Once again, HIRE successfully recovers the six underlying reference cell types and estimates the cellular compositions well (see Methods).

### Real data analysis

HIRE also provides more insights into real data than the previous studies. The rheumatoid arthritis (RA) dataset [3] contains methylation profiles collected from the whole blood of 354 RA patients and 335 normal participants. Besides the RA status, other attributes such as gender, smoking history, age, and batch information are also available. We first corrected the batch effects and then applied HIRE to the dataset (Methods). Fig. 3a displays the p-values regarding the association to the RA status for each CpG site in each cell type, where HIRE selected six cell types (Supplementary Fig. 63a), consistent with the cell type number in [13]. Despite any potential batch effects and biological variability, three out of the six cell types can be matched to known blood cell references—cell type 1 to CD4+ T cells, cell type 2 to neutrophils, cell type 4 also to neutrophils, and the remaining three cell types cannot be aligned to the references (Methods and Supplementary Fig. 64). HIRE detected 63 risk-CpG sites in cell type 3—the largest number of associations across all of the cell types—but 0 risk-CpG sites in cell type 1 (Supplementary Table 1). Therefore, the disease status affected some but not necessarily all of the cell types. Note that the significant CpG site cg06373940 called by HIRE is located on gene ERCC3. The level of ERCC3’s corresponding protein has been reported to increase in RA synovium [25]. Moreover, we found five CpG sites to be significantly associated with smoking history (Supplementary Fig. 65 and Supplementary Table 2). One of them is cg05575921, which has recently been linked to smoking in two other independent studies with blood samples [26, 27]. However, these findings were missed by the association detection at the aggregated level in the previous analyses of the same dataset [11, 13]. The p-value density plots and Q-Q plots for the commonly used methods are also displayed in Figs. 3c-n; they present patterns similar to those observed in the simulation study except that there is obvious overcorrection by ReFACTor.

**Figure 3:**
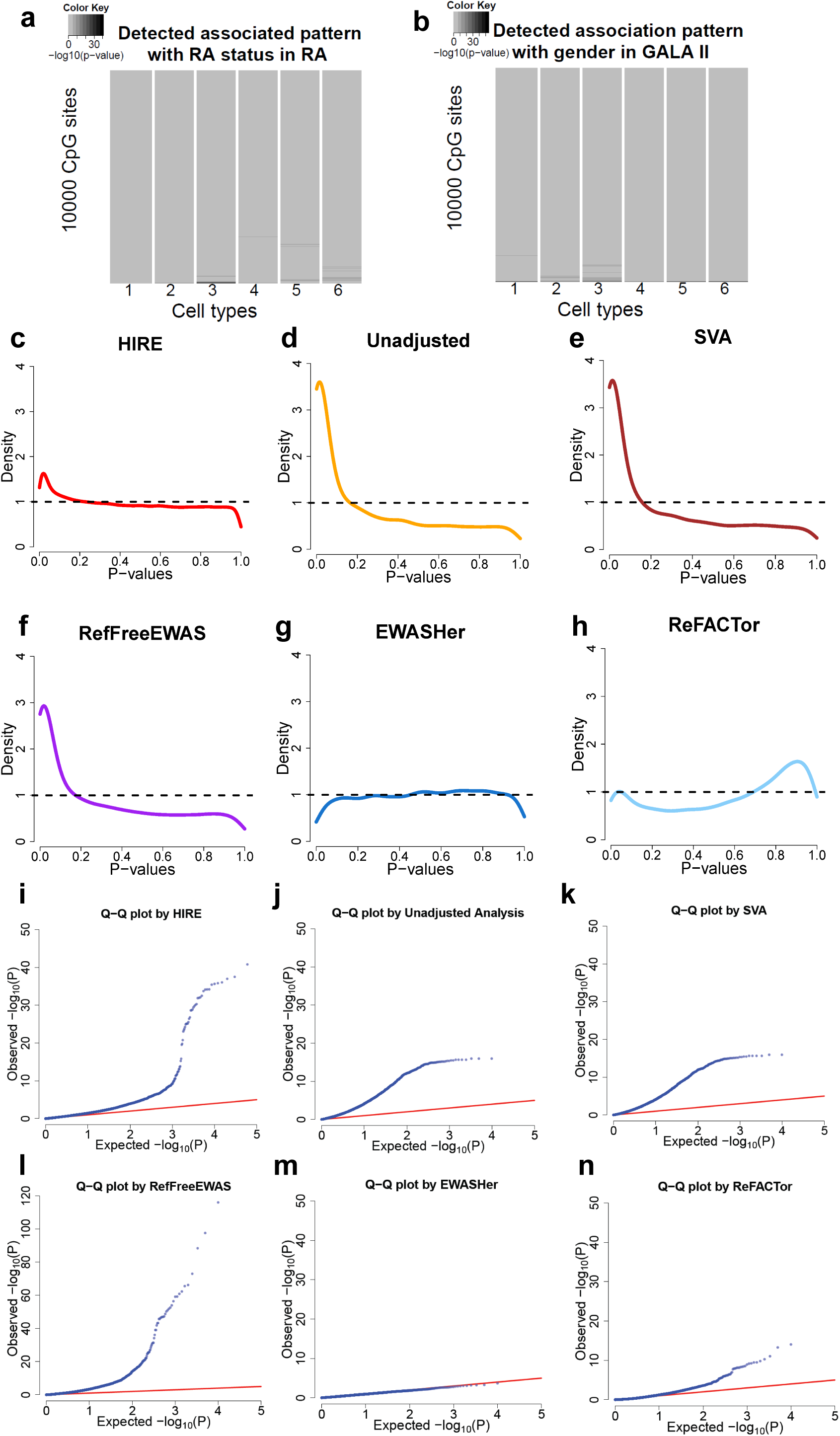
The application of HIRE and the commonly used methods to two real methylation datasets: RA and GALA II. (**a**) The cell-type-specific association pattern with RA status detected by HIRE in the RA dataset. The darkness represents the *-* log_10_(*p - value*). (**b**) The cell-type-specific association pattern with gender detected by HIRE in the GALA II dataset. The darkness represents the *-* log_10_(*p - value*). (**c-h**) The p-value density plots for association with RA status in the RA dataset for (**c**) HIRE, (**d**) unadjusted analysis, (**e**) SVA, (**f**) RefFreeEWAS, (**g**) EWASHer, and (**h**) ReFACTor. (**i-n**) The Q-Q plots for association with RA status in the RA dataset for (**i**) HIRE, (**j**) unadjusted analysis, (**k**) SVA, (**l**) RefFreeEWAS, (**m**) EWASHer, and (**n**) ReFACTor.

The high resolution provided by HIRE makes it a powerful tool for EWAS studies. Rahmani et al. analyzed the GALA II blood methylation dataset [28] with ReFACTor [13]. The dataset consists of 573 samples collected from a pediatric Latino population. Each sample has the gender information and belongs to one of the following four populations: “Mexican,”“Mixed Latino,”“Puerto Rican,” and “Other Latino.” We applied HIRE to the dataset to investigate whether there were any cell-type-specific CpG sites associated with gender and ethnicity. We created three dummy variables to represent the four ethnic groups. By taking indicators of ethnicity as phenotypes in the model, HIRE automatically accounts for the population differences in cell compositions and cell-type-specific methylation levels simultaneously. HIRE correctly selected the number of cell types as six as reported in [13] (Supplementary Fig. 63b). According to cell-type alignment, cell types 1 and 5 can be annotated as CD4+ T cells; cell types 2, 3, and 4 belong to neutrophils; and cell type 6 was annotated as CD56+ natural killer cell (CD56+ NK) using the references (Supplementary Fig. 66). HIRE found 1936 CpG sites to be associated with ethnicity across all of the cell types (Supplementary Fig. 67) and identified 14, 52, 155, 15, 18, and 14 risk-CpG sites of the gender in cell types 1-6, respectively (Fig. 3b). The gene set enrichment analysis showed that the genes that harbored the risk-CpG sites for gender were significantly enriched in seven canonical pathways (Supplementary Table 3), of which PID CMYB PATHWAY was ranked the highest. The transcription factor *c - MY B* in PID CMYB PATHWAY enhances the progression of breast cancer [29]. Therefore, the different occurrence rates of breast cancers between males and females may be linked to the differences at the epigenome level. In comparison, only one pathway was found to be enriched with the genes hosting the risk-CpG sites claimed by ReFACTor at the aggregated level (Supplementary Table 4). All of these observations highlight the importance of the finer-scale resolutions of HIRE.

## Discussion

In reality, the phenotype may affect a risk-CpG site in some but not all of the cell types. As far as we know, HIRE is the first tool to detect the cell-type-specific association pattern with each phenotype for EWAS. The identification of cell-type-specific risk-CpG sites will help epigenetic therapies to target the affected cell types more effectively.

Statistically, instead of assuming a fixed reference methylomes for all the samples as the existing methods do [9, 13, 16], HIRE allows each sample’s cell-type-specific methylation profiles to depend on its phenotypes. Consequently, HIRE correctly models the multiplicative effects of the cellular compositions on the observed methylation levels, whereas the existing approaches all misspecify the cellular compositions as additive effects (Methods). As a result, HIRE enables the detection of cell-type-specific risk-CpG sites infeasible by existing state-of-the-art methods. As a byproduct, HIRE also improves the statistical power of the association detection at the aggregated level over existing state-of-the-art methods. Computationally, the time complexity of one iteration by HIRE is *O*(*nmKp* + *nK*^3^), thus providing a fast convergence when *K* is moderate. The statistical and computational advantages equip HIRE to scale up for large-cohort EWAS.

So far, in the EWAS community, there is no gold-standard dataset to compare different methods of detecting associations. Ideally, we would like to have epigenetic spike-in experiments where purified cell types are first isolated, then CpG-sites are epigenetically edited on a per cell type basis, and finally cell types are mixed in predetermined proportions. Given such experiments, we know the underlying truth about which CpGs are differentially methylated in each cell type and the cell mixing proportions for each sample. However, biotechnologies for epigenetic editing, such as CRISPR-Cas, are still not mature at this stage, with many off-targets modifications [30]. Therefore, most computational EWAS papers refer to numerical simulation studies rather than to experimental studies when evaluating the performance of their algorithms [12, 13]. Here, we follow previous comparative studies and design our simulation studies to serve as the computational counterpart of experimental spike-in studies. With the rapid advancement in epigenetic editing, we hope the community can devote more effort in the near future to creating a gold-standard dataset, such as those generated in the early years for gene expression microarray studies [31].

The beta-values representing methylation levels are always between zero and one. As previous approaches to EWAS often assume normal distribution for the beta-values and show good performances in real applications [9, 13], in HIRE, we also assume that the beta-values follow normal distributions. Consequently, there is a chance that the fitted methylation level is out of the range of [0, 1]. Nevertheless, we do in fact constrain the baseline methylation profiles *µ*_*jk*_s to the closed interval [0, 1] and force the cellular compositions *p*_*ki*_s to be non-negative and sum up to one: 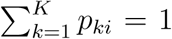. As a result, as the phenotypes have no effect on most CpG sites, most observations, *O*_*ji*_s, have their means 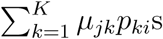 in [0, 1]. In fact, for both the RA dataset and the GALA II dataset, more than 99.99% of the fitted methylation values Ô_jis_ based on HIRE estimates are between zero and one. Therefore, the normal assumption fits the data reasonably well and does not have a large effect on the performance of HIRE.

One major issue for all of the cell-type deconvolution methods is that deconvolution cannot be achieved if there is no variation of cellular compositions among samples. For example, assuming that the samples are mixtures of two cell types and **p**_*i*_ = **p** for all of the samples, then the observed methylation profile **O**_*i*_ equals **u**_*i*1_*p*_1_ + **u**_*i*2_*p*_2_ = (**u**_*i*1_ + *p*_2_*C*)*p*_1_ + (**u**_*i*2_ *- p*_1_*C*)*p*_2_:= Ũi1*p*_1_ + Ũi2*p*_2_ for any constant *C*. As a result, **u**_*i*1_ and **u**_*i*2_ are not estimable. As HIRE is also a deconvolution-based method, it suffers from the same problem. However, for tissues with large cellular heterogeneity such as blood, deconvolution-based methods are applicable, and HIRE can accurately estimate cellular compositions.

HIRE requires a moderate sample size to obtain precise estimates as HIRE needs to learn (1 + 2*K* + *qK*)*m* + (*K -* 1)*n* parameters with a total of *mn* observed values. Our simulation studies show that with 180 samples, HIRE performs very well at the aggregated level (Table 1). Suppose the sample sizes drop below 150, say to 120, HIRE can still control the FPR well but begins to lose power (Supplementary Table 5). Like the two datasets analyzed in the real application, a typical sample size for a current EWAS is over 500, thus guaranteeing a high TPR for HIRE. Given the decreasing cost of EWAS, we recommend that researchers collect at least 200 samples for their studies for association detection at the aggregated level and 600 samples for identifying cell-type-specific risk-CpG sites. A larger sample sizes can further boost the power.

With the popularity of EWAS, we believe that HIRE will be widely applied, and we hope that HIRE can motivate more researchers to mine out finer-scale results from EWAS.

## Methods

### Multiplicative effects of cellular composition on methylation

In this section, we illustrate that the effects of the cell-type composition are actually *multiplicative*. Let us assume that the beta-values representing the methylation levels are observed across *m* CpG sites for *n* samples. As the measured sample is composed of cells from different cell types, the observed beta-value is a weighted average of the mean methylation levels of distinct cell types, and the weights correspond to the proportions of each cell type. Let *O*_*ji*_ denote the measurement at CpG site *j* for sample *i*. If we assume that there exist *K* cell types in all of the samples and the mean methylation level for CpG site *j* in cell type *k* is *µ*_*jk*_, then

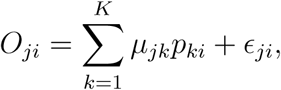

where *p*_*ki*_ is the proportion of cell type *k* in sample *i* with a natural constraint 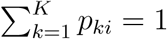 and *E*_*ji*_ is a random error.

Let us consider a case-control EWAS. Without loss of generality, we assume that CpG site *j* is differentially methylated between cases and controls in cell type 1 with a mean shift *δ*_*j*1_ and it is not differentially methylated in the rest of the cell types. As a result, for case samples,

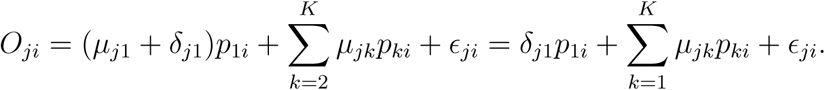

Subsequently, if we use *Z*_*i*_ to indicate the case-control status of sample *i*, the observed methylation level becomes

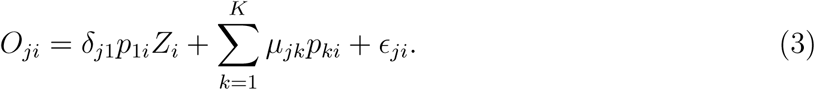

Therefore, the proportions of cell type 1—*p*_1*i*_, *i* = 1, *…, n*—have multiplicative effects rather than additive effects on the mean difference between the case and the control samples.

The existing methods, either estimating the cell type proportions explicitly or approximating them with surrogate variables implicitly, add the estimated proportions and the case-control indicator *Z*_*i*_ as the covariates to the regression as follows:

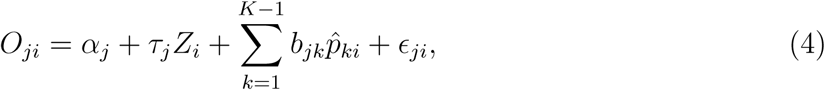

where *b*_*jk*_s are the regression coefficients. As a result, CpG site *j* is called differentially methylated on the basis of hypothesis test for *τ*_*j*_ = 0. In general, *τ*_*j*_ in Equation (4) is not equal to *δ*_*j*1_ in Equation (3). Please see the Supplementary Note for a numerical example. Moreover, testing for *τ*_*j*_ = 0 loses the information about the cell type in which the CpG site *j* may be at risk. Accounting for the multiplicative effects, we propose the HIRE model that conserves the individual cell-type level information, which is introduced in the next section.

### The HIRE model

HIRE employs a hierarchical model to closely follow the data generation process for the EWAS data. To begin with, we assume that the baseline methylation level for CpG site *j* in cell type *k* is *µ*_*jk*_. For sample *i* with phenotypes **x**_*i*_ = (*x*_*i*1_*,…, x*_*iq*_), the mean methylation value for CpG site *j* in cell type *k* is assumed to be 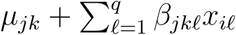 In other words, the phenotypes have linear effects where *β*_*jk*_ characterizes the influence of phenotype *𝓁* on CpG site *j* in cell type *k*. Let *u*_*ijk*_ represent the signal from CpG site *j* in cell type *k* for sample *i* with **x**_*i*_. We assume that *u*_*ijk*_ is normally distributed with mean 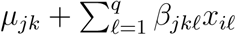 and standard deviation *σ*_*jk*_,

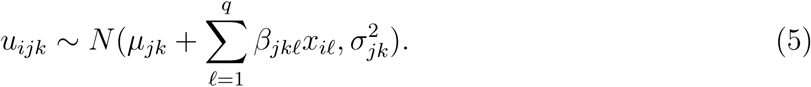

After *u*_*ijk*_s are generated for all of the *K* cell types, the observed methylation value *O*_*ji*_ is sampled as follows:

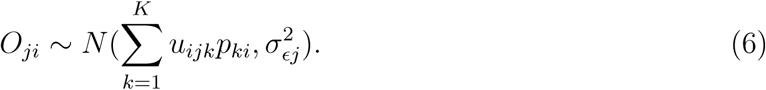

Collectively, **O** = *{O*_*ji*_ : 1 *≤ j ≤ m,* 1 *≤ i ≤ n}* denote the observed data; **u** = *{*(*u*_*ij*1_*,…, u*_*ijK*_)^*T*^ : 1 *≤ i ≤ n,* 1 *≤ j ≤ m}* are the missing data; and ***µ***_*j*_ = (*µ*_*j*1_*,…, µ*_*jK*_)^*T*^, 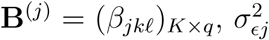 the diagonal matrix 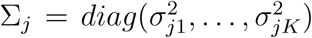 for *j* = 1*,…, m*, and **p**_*i*_ = (*p*_1*i*_*,…, p*_*Ki*_)^*T*^ for *i* = 1*,…, n* are the parameters. With 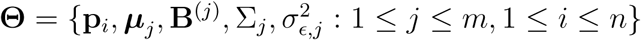 the complete data log-likelihood function, *l*_*c*_, can be expressed as follows:

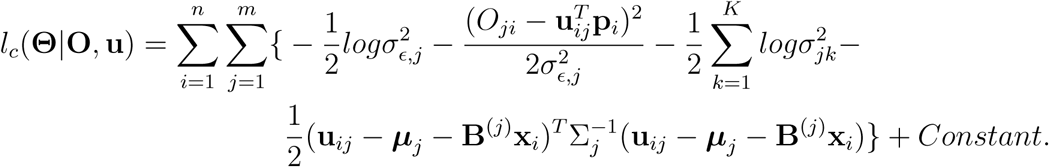

Accordingly, we develop a generalized expectation-maximization algorithm [32] to estimate the parameters. In expectation-maximization algorithm, a good initialization can lead to faster convergence than random starts. We adopt the cellular composition estimations from the methylation matrix decomposition algorithm [16] with slight modifications as the initializations. The initial values for the baseline methylation profiles *µ*_*jk*_ are accordingly estimated by simple linear regressions. As the number of risk-CpG sites is often small, all of the phenotype effects *β*_*jk*_ are set to zero at the beginning. For the standard deviations, the initial values are randomly sampled from inverse gamma distributions with small means. We choose the number of cell types *K* by using a variant of the penalized Bayesian information criterion (pBIC) [20] (see details in Supplementary Note).

For each phenotype *𝓁*, we can conduct the hypothesis test *H*_0_ : *β*_*jk*_ = 0 versus *H*_1_ : *β*_*jk*_ *≠* 0 for any cell type *k* and any CpG site *j*. Combining Equations (5) and (6), we obtain the following equations:

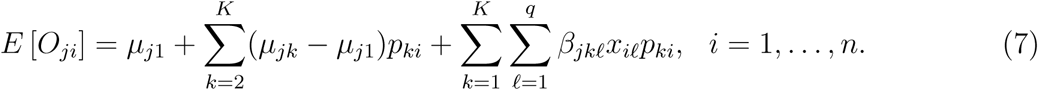

Subsequently, we can take (*O*_*j*1_*,…, O*_*jn*_) as the response vector and concatenate **1**_*n*_, (*p*_*k*1_*,…, p*_*kn*_) (*k* = 2*,…, K*) and (*x*_1_ *p*_*k*1_*,…, x*_*n*_ *p*_*kn*_) (*𝓁* = 1*,…, q*; *k* = 1*,…, K*) to a *n ×* (*p* + 1) *K* design matrix in the linear regression. We plug in the estimated cellular compositions *p*_*ki*_ and conduct the hypothesis test for *β*_*jk*_ = 0 using the t-tests in the linear models. We claim that CpG site *j* has association with phenotype *𝓁* at the aggregated level if phenotype *𝓁* affects CpG site *j* in at least one of the *K* cell types. Note that in the regression we incorporate the estimated cellular compositions into the linear model as multiplicative effects rather than additive effects.

More technical details of the method and the algorithm are available in the Supplementary Note.

### Data simulation

We compared the performance of HIRE with five previous methods— unadjusted analysis, SVA, RefFreeEWAS, EWASHer, and ReFACTor—in 18 simulation settings. We set the sample size *n* to 180, 300, and 600 and let the underlying cell type number *K* be 3, 5, and 7. For each pair of (*n, K*), we investigated the “*true null* ” case and the “*true alternative*” case. As a result, we have in total 3 (the number of sample sizes) *×* 3 (the number of cell types) *×* 2 (the “true null” case and the “true alternative” case) = 18 simulation settings. For each setting, we considered 10,000 CpG sites and simultaneously accounted for the following factors.

#### Cell lineage

We first constructed the baseline methylation matrix ***µ*** = (*µ*_*jk*_)_*m×K*_, where each column corresponds to the baseline methylation levels of a cell type. To mimic the phenomenon that cell types from the same lineage have similar methylation profiles, we assumed that *K*_*sim*_ out of the total *K* cell types were similar. Specifically, without loss of generality, we assumed that the first *K*_*sim*_ cell types were from the same cell lineage and the rest *K - K*_*sim*_ cell types are irrelevant to one another. We set *K*_*sim*_ to 2, 2, and 3 for *K* = 3, 5, and 7, respectively. We generated *µ*_*jk*_ for cell types *k* = 1, *K*_*sim*_ + 1*,…, K* from the beta distribution *beta*(3, 6) on each CpG site *j* independently. For each of the remaining cell types *k*^*l*^ = 2*,…, K*_*sim*_, we randomly selected 20% of the CpG sites and drew their *µ*_*jk*_*t* s independently from *beta*(3, 6); and for the remaining 80% of CpG sites, we let their *µ*_*jk*_*t* be *µ*_*j*1_ plus a very small randomness, thus inducing the similarities among cell types 1 to *K*_*sim*_.

#### Discrete and continuous phenotypes

We further generated a discrete and a continuous phenotype **x** = (**x**_1_, **x**_2_)^*T*^ for each individual *i* (*i* = 1*,…, n*). We let the first *n/*3 individuals be the control samples with *x*_*i*1_ = 0 for *i* = 1*,…, n/*3 and the remaining 2*n/*3 individuals serve as cases with *x*_*i*1_ = 1 for *i* = *n/*3 + 1*,…, n*. The continuous phenotypes **x**_2_ = (*x*_12_*,…, x*_*i*2_*,…, x*_*n*2_)^*T*^ were independently drawn from a *Unif* (20, 50) to act as age.

*Phenotype effects with different magnitudes and directions*. Subsequently, we simulated the phenotype effect *β*_*jk*_ of each phenotype *𝓁* on CpG site *j* in cell type *k*. For the “true null” cases, all of the *β*_*jk*_ s are zero. For a “true alternative” setting, we set nonzero phenotype effects as follows.

For phenotype 1—the case/control status, we let it affect the first 10 CpG sites in all of the cell types: *β*_*jk*1_ *≠* 0 for *j* = 1*,…,* 10 and *k* = 1*,…, K*. Then, we assumed that the next 10 CpG sites were influenced by the disease status in the first *K*_*sim*_ cell types which come from the same lineage but not the other cell types: *β*_*jk*1_ *≠* 0 (*k* = 1*,…, K*_*sim*_) and *β*_*jk*1_ = 0 (*k* = *K*_*sim*_ + 1*,…, K*) for any *j* = 11, *…,* 20. Furthermore, for cell type *k ∈ {K*_*sim*_ + 1*,…, K}*, we let the disease status affect CpG sites *j* = 20 + 10(*k - K*_*sim*_ *-* 1) + 1*,…,* 20 + 10(*k - K*_*sim*_) only in cell type *k*. We generated the cell-type-specific effects of age in a similar fashion for CpG site loci 21 to 40 + 10(*K - K*_*sim*_).

For each nonzero *β*_*jk*1_, we let *β*_*jk*1_ = *r*_*jk*_ *· ω*_*jk*_, where *ω*_*jk*_ *∼ Unif* (0.07, 0.15) and *r*_*jk*_ takes values at 1 and *-*1 with equal probabilities. Thus, *β*_*jk*1_s can have both positive and negative effects. In the same spirit, we generated nonzero *β*_*jk*1_s with 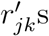 and 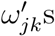 where 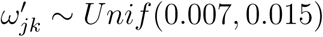.

#### The association between phenotypes and cellular compositions

Notice that the phenotypes may be associated with the cellular compositions. Therefore, when *K* = 3, we drew **p**_*i*_ = (*p*_1*i*_*,…, p*_*Ki*_) from a Dirichlet distribution *Dir*(4, 4, 2 + 0.1*x*_*i*2_) if sample *i* is a control and **p**_*i*_ *∼ Dir*(4, 4, 5 + 0.1*x*_*i*2_) if it is a case; when *K* = 5, we let **p**_*i*_ *∼ Dir*(3, 3, 3, 3, 2 + 0.1*x*_*i*2_) for a control sample and **p**_*i*_ *∼ Dir*(3, 3, 3, 3, 5 + 0.1*x*_*i*2_) for a case sample; and when *K* = 7, we sampled **p**_*i*_ *∼ Dir*(1, 3, 3, 3, 2, 2, 2 + 0.1*x*_*i*2_) for controls and **p**_*i*_ *∼ Dir*(1, 3, 3, 3, 2, 2, 5 + 0.1*x*_*i*2_) for cases.

Finally, we generated the observed value *O*_*ji*_ for CpG site *j* of sample *i* as follows: sample *u*_*ijk*_ from *N* (*µ*_*jk*_ + *β*_*jk*1_*x*_*i*1_ + *β*_*jk*2_*x*_*i*2_, 0.01^2^) for *k* = 1*,…, K*; and sample *O*_*ji*_ from 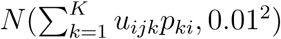 In case *O*_*ji*_ is out of the interval (0, 1), we truncate it to zero if *O*_*ji*_ is lower than zero and to one if *O*_*ji*_ is greater than one.

### A semi-simulated dataset including samples with known cell mix proportions

The GEO dataset GSE110554 [24] contains purified cell-type-specific methylation profiles for six cell types: neutrophils, monocytes, B cells, CD4+ T, CD8+ T, and NK. Moreover, GSE110554 includes mixed samples, whose methylation signals were aggregated from the six cell types with pre-determined cell mix proportions. Therefore, because of the known cell type and cellular proportion information, GSE110554 is an ideal dataset with which to test HIRE’s performance.

In GSE110554, the number of mixed samples is much smaller than the typical size of an EWAS and, as discussed in the manuscript, HIRE usually requires hundreds of samples to obtain accurate and stable results. Therefore, to increase the sample size, we first generated a simulated methylation dataset with 600 samples using the purified methylation profiles. We focused on 10k CpG sites including the 450 IDOL CpG sites, which were previously identified as the optimal library of CpG sites for estimating leukocyte subtype proportions [24], and another 9550 CpG sites whose methylation values across the purified cell types were in the range of [0.2, 0.8] and had large standard deviations [11]. Subsequently, we combined the 600 samples and 6 mixed samples (generated by method A) [24] available in GSE110554 to compose a semi-simulated dataset.

After applying HIRE to the semi-simulated data, we annotated the estimated cell types based on the methylation profiles from GSE110554. Supplementary Fig. 69 shows the heatmap for the Pearson correlation matrix between inferred cell types and the underlying truth. The correlation signals on the diagonal are the strongest in each row. HIRE successfully recovers the six underlying cell types. We also compared the estimated cellular compositions with the underlying true proportions for the 6 mixed samples. In Supplementary Fig. 70, each panel displays the scatter plot between the cellular proportion estimates and the true mix proportions for a given cell type, all of which indicate that HIRE obtains good estimates for cellular compositions.

### Cell type matching protocol

Assume that we have the reference methylation profiles for the *H* annotated cell types. We denote the methylation profile for cell type *h* as ***ϕ***_*h*_ = (*ϕ*_1*h*_*,…, ϕ*_*mh*_). We aim to annotate ***µ***_*k*_ using the references. Following [33], first, we calculate the cosine similarity, the Pearson correlation, and the Spearman correlation between ***µ***_*k*_ and ***ϕ***_*h*_ for each cell type *h ∈ {*1*,…, H}*. Notice that the three similarity measures are between *-*1 and 1 and a high positive value indicates a high similarity between two vectors. Second, for each similarity measure *𝓁* (*𝓁* = 1, 2, 3), we identify the cell type *h* that has the maximal similarity degree with ***µ***_*k.*_ If at least two out of the three similarity measures identify the same reference cell type 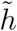 and their corresponding similarity values are greater than 0.5, then we annotate ***µ***_*k*_ with the reference cell type *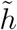*. Otherwise, ***µ***_*k*_ is believed to belong to a “new” cell type that is not included in the references. We repeat the above process for each methylation profile ***µ***_*k*_ estimated from HIRE.

### The blood cell references

The two real data sets analyzed in our applications were obtained from whole blood. Therefore, we prepared the references from a whole blood methylation study [34] with GEO accession code GSE35069. The study collected seven isolated blood cell subpopulations—CD4+ T cells, CD8+ T cells, CD14+ monocytes, CD19+ B cells, CD56+ NK cells, neutrophils, and eosinophils—for six individuals. Accordingly, we define the reference profile ***ϕ***_*h*_ for cell type *h* as the average methylation profile of these individuals, i.e 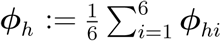

### Data preprocessing

The RA dataset is publicly available in GEO with accession number GSE42861. The dataset measures the methylation levels of the whole blood. The methylation data have been normalized by Illumina’s control probe scaling procedure (see Liu et al. [3] “Illumina 450K microarray data preprocessing” section for details). There are in total 689 samples. For each sample, his or her RA status, age, gender, smoking history, and batch information are also available. We removed two samples “GSM1051535” and “GSM1051691” as their smoking information is missing. CpG sites with a high methylation mean (*>* 0.8) and a low methylation mean (*<* 0.2) were discarded [11, 13]. We adjusted the data for batch effects using COMBAT [35]. The correction process was justified as we did not observe a high co-linearity between the RA status and the batches (Supplementary Fig. 68). Subsequently, the 10,000 most variable CpG sites were kept. For the RA status, we denoted the RA patient by 1 and the normal by 0; we represented male with 1 and female with 0; for the smoking history, we used (0, 0, 0) to refer to “never,” (1, 0, 0) to “ex,” (0, 1, 0) to “current,” and (0, 0, 1) to “occasional” smokers.

We downloaded the GALA II dataset from Gene Expression Omnibus (GEO) with accession number GSE77716. The dataset contains the whole blood DNA methylation beta-values from 573 samples. The beta-values have been normalized by SWAN [36] and corrected for batch effects by COMBAT [35]. There are two types of covariates, the gender and the ethnicity. The ethnicity includes “Mexican,” “Mixed Latino,” “Puerto Rican,” and “Other Latino.” Out of the 573 samples, one sample “GSM2057284” has no gender information, so we removed it. As suggested by previous studies [11, 13], CpG sites with a mean methylation value of less than 0.2 or higher than 0.8 were filtered out. Among the remaining CpG sites, we selected the 10,000 most variable CpG sites. For gender, we denoted male by 1 and female by 0. For the ethnicity variables, we used three dummy variables to represent the four ethnicity categories. In particular, (0, 0, 0), (1, 0, 0), (0, 1, 0), and (0, 0, 1) corresponded to “Mexican,” “Mixed Latino,” “Puerto Rican,” and “Other Latino,” respectively.

For ReFACTor and EWASHer, according to their rules, we first filtered out CpG sites that were consistently hypomethylated or consistently hypermethylated and then regressed out the known covariates. We finally used the residuals to perform their analysis. Note that in their software these steps are processed automatically. For RefFreeEWAS, SVA, and the unadjusted analysis, the phenotypes and the covariates were regarded as the fixed effects in the regression model. In detail, for ReFACTor, in both GALA II and RA datasets, the cell type number “K” was specified to be six, which was the same as in their paper [13]. For RefFreeEWAS, we fixed the dimensionality of latent space “d” at six in the real data. For SVA, we also fixed the number of surrogate variables to six.

The gene enrichment analysis was carried out on the broad institute website http://software.broadinstitute.org/gsea/msigdb/annotate.jsp. The canonical pathways were selected as the basis gene sets, and only pathways with a false discovery rate of less than 0.05 were reported.

### Code availability

The R package to implement HIRE is available on GitHub: https://github.com/XiangyuLuo/HIREewas.

### Data availability

The RA whole blood methylation dataset is available in the Gene Expression Omnibus (GEO) with the accession number GSE42861. The GALA II whole blood methylation dataset can be downloaded from GEO with the accession number GSE77716. The accession number for the blood cell references is GSE35069. The purified methylation data and mixed samples used to generate the semi-simulated dataset are from GSE110554.

## CONTRIBUTIONS

YY.W. and C.Y. conceived the study. XY.L. and YY.W. developed the method. XY.L. implemented the algorithm and prepared the software package. XY.L. and C.Y. analyzed the data. XY.L., YY.W. and C.Y. wrote the paper.

## References

[1] Vardhman K Rakyan, Thomas A Down, David J Balding, and Stephan Beck. Epigenome-wide association studies for common human diseases. Nature Reviews Genetics, 12(8):529–541., 2011.

[2] Mukesh Verma. Epigenome-wide association studies (EWAS) in cancer. Current Genomics, 13(4):308–313., 2012.

[3] Y-un Liu, Martin J Aryee, Leonid Padyukov, M Daniele Fallin, Espen Hesselberg, Arni Runarsson, Lovisa Reinius, Nathalie Acevedo, Margaret Taub, Marcus Ronninger, et al. Epigenome-wide association data implicate DNA methylation as an intermediary of genetic risk in rheumatoid arthritis. Nature Biotechnology, 31(2):142–147., 2013.

[4] Xu Gao, Min Jia, Yan Zhang, Lutz Philipp Breitling, and Hermann Brenner. DNA methylation changes of whole blood cells in response to active smoking exposure in adults: a systematic review of DNA methylation studies. Clinical Epigenetics, 7(1):113, 2015.

[5] Roby Joehanes, Allan C Just, Riccardo E Marioni, Luke C Pilling, Lindsay M Reynolds, Pooja R Mandaviya, Weihua Guan, Tao Xu, Cathy E Elks, Stella Aslibekyan, et al. Epigenetic signatures of cigarette smoking. Circulation: Genomic and Precision Medicine, 9(5):436–447., 2016.

[6] Simone Wahl, Alexander Drong, Benjamin Lehne, Marie Loh, William R Scott, Sonja Kunze, Pei-Chien Tsai, Janina S Ried, Weihua Zhang, Youwen Yang, et al. Epigenome-wide association study of body mass index, and the adverse outcomes of adiposity. Nature, 541(7635):81–86., 2017.

[7] Andrew E Teschendorff, Usha Menon, Aleksandra Gentry-Maharaj, Susan J Ramus, Daniel J Weisenberger, Hui Shen, Mihaela Campan, Houtan Noushmehr, Christopher G Bell, A Peter Maxwell, et al. Age-dependent DNA methylation of genes that are suppressed in stem cells is a hallmark of cancer. Genome Research, 20(4):440–446., 2010.

[8] Steve Horvath. DNA methylation age of human tissues and cell types. Genome Biology, 14(10):3156, 2013.

[9] Eugene Andres Houseman, William P Accomando, Devin C Koestler, Brock C Christensen, Carmen J Marsit, Heather H Nelson, John K Wiencke, and Karl T Kelsey. DNA methylation arrays as surrogate measures of cell mixture distribution. BMC Bioinformatics, 13(1):86, 2012.

[10] Andrew E Jaffe and Rafael A Irizarry. Accounting for cellular heterogeneity is critical in epigenome-wide association studies. Genome Biology, 15(2):R31, 2014.

[11] James Zou, Christoph Lippert, David Heckerman, Martin Aryee, and Jennifer Listgarten. Epigenome-wide association studies without the need for cell-type composition. Nature Methods, 11(3):309–311., 2014.

[12] Kevin McGregor, Sasha Bernatsky, Ines Colmegna, Marie Hudson, Tomi Pastinen, Aurélie Labbe, and Celia MT Greenwood. An evaluation of methods correcting for cell-type heterogeneity in DNA methylation studies. Genome Biology, 17(1):84, 2016.

[13] Elior Rahmani, Noah Zaitlen, Yael Baran, Celeste Eng, Donglei Hu, Joshua Galanter, Sam Oh, Esteban G Burchard, Eleazar Eskin, James Zou, et al. Sparse PCA corrects for cell type heterogeneity in epigenome-wide association studies. Nature Methods, 13(5):443–445., 2016.

[14] Andrew E Teschendorff and Caroline L Relton. Statistical and integrative system-level analysis of DNA methylation data. Nature Reviews Genetics, 2017.

[15] William P Accomando, John K Wiencke, E Andres Houseman, Heather H Nelson, and Karl T Kelsey. Quantitative reconstruction of leukocyte subsets using DNA methylation. Genome Biology, 15(3):R50, 2014.

[16] Eugene Andres Houseman, Molly L Kile, David C Christiani, Tan A Ince, Karl T Kelsey, and Carmen J Marsit. Reference-free deconvolution of DNA methylation data and mediation by cell composition effects. BMC Bioinformatics, 17(1):259, 2016.

[17] Xiaoqi Zheng, Naiqian Zhang, Hua-Jun Wu, and Hao Wu. Estimating and accounting for tumor purity in the analysis of DNA methylation data from cancer studies,. Genome Biology, 18(1):17, 2017.

[18] Jeffrey T Leek and John D Storey. Capturing heterogeneity in gene expression studies by surrogate variable analysis. PLoS Genetics, 3(9):e161, 2007.

[19] Eugene Andres Houseman, John Molitor, and Carmen J Marsit. Reference-free cell mixture adjustments in analysis of DNA methylation data. Bioinformatics, 30(10):1431–1439., 2014.

[20] Wei Pan and Xiaotong Shen. Penalized model-based clustering with application to variable selection. Journal of Machine Learning Research, 8(May):1145–1164., 2007.

[21] Shijie C Zheng, Stephan Beck, Andrew E Jaffe, Devin C Koestler, Kasper D Hansen, Andres E Houseman, Rafael A Irizarry, and Andrew E Teschendorff. Correcting for cell-type heterogeneity in epigenome-wide association studies: revisiting previous analyses. Nature Methods, 14(3):216–217., 2017.

[22] Elior Rahmani, Noah Zaitlen, Yael Baran, Celeste Eng, Donglei Hu, Joshua Galanter, Sam Oh, Esteban G Burchard, Eleazar Eskin, James Zou, et al. Correcting for cell-type heterogeneity in DNA methylation: a comprehensive evaluation. Nature Methods, 14(3):218–219., 2017.

[23] John D Storey and Robert Tibshirani. Statistical significance for genomewide studies. Proceedings of the National Academy of Sciences, 100(16):9440–9445., 2003.

[24] Lucas A Salas, Devin C Koestler, Rondi A Butler, Helen M Hansen, John K Wiencke, Karl T Kelsey, and Brock C Christensen. An optimized library for reference-based deconvolution of whole-blood biospecimens assayed using the illumina humanmethylationepic beadarray. Genome Biology, 19(1):64, 2018.

[25] E Neumann, F Kullmann, M Judex, HP Jüsten, D Wessinghage, S Gay, J Schölmerich, and U Müller-Ladner. Identification of differentially expressed genes in rheumatoid arthritis by a combination of complementary DNA array and rna arbitrarily primed-polymerase chain reaction. Arthritis & Rheumatology, 46(1):52–63., 2002.

[26] Francesca Fasanelli, Laura Baglietto, Erica Ponzi, Florence Guida, Gianluca Campanella, Mattias Johansson, Kjell Grankvist, Mikael Johansson, Manuela Bianca Assumma, Alessio Naccarati, et al. Hypomethylation of smoking-related genes is associated with future lung cancer in four prospective cohorts. Nature Communications, 6:10192, 2015.

[27] Srikant Ambatipudi, Cyrille Cuenin, Hector Hernandez-Vargas, Akram Ghantous, Florence Le Calvez-Kelm, Rudolf Kaaks, Myrto Barrdahl, Heiner Boeing, Krasimira Aleksandrova, Antonia Trichopoulou, et al. Tobacco smoking-associated genome-wide DNA methylation changes in the epic study. Epigenomics, 8(5):599–618., 2016.

[28] Maria Pino-Yanes, Neeta Thakur, Christopher R Gignoux, Joshua M Galanter, Lindsey A Roth, Celeste Eng, Katherine K Nishimura, Sam S Oh, Hita Vora, Scott Huntsman, et al. Genetic ancestry influences asthma susceptibility and lung function among latinos. Journal of Allergy and Clinical Immunology, 135(1):228–235., 2015.

[29] Yihao Li, Ke Jin, Gabi W van Pelt, Hans van Dam, Xiao Yu, Wilma E Mesker, Peter ten Dijke, Fangfang Zhou, and Long Zhang. c-Myb enhances breast cancer invasion and metastasis through the wnt/*β*-catenin/axin2 pathway. Cancer Research, 76(11):3364–3375., 2016.

[30] Xiao-Hui Zhang, Louis Y Tee, Xiao-Gang Wang, Qun-Shan Huang, and Shi-Hua Yang. Off-target effects in CRISPR/Cas9-mediated genome engineering. Molecular Therapy-Nucleic Acids, 4, 2015.

[31] Benjamin M Bolstad, Rafael A Irizarry, Magnus Åstrand, and Terence P. Speed. A comparison of normalization methods for high density oligonucleotide array data based on variance and bias. Bioinformatics, 19(2):185–193., 2003.

[32] Arthur P Dempster, Nan M Laird, and Donald B Rubin. Maximum likelihood from incomplete data via the EM algorithm. Journal of the Royal Statistical Society. Series B (Methodological), pages 1–38., 1977.

[33] Vladimir Yu Kiselev,Andrew Yiu, and Martin Hemberg. scmap: projection of single-cell RNA-seq data across data sets. Nature methods, 15(5):359, 2018.

[34] Lovisa E Reinius, Nathalie Acevedo, Maaike Joerink, Göran Pershagen, Sven-Erik Dahlén, Dario Greco, Cilla Söderhäll, Annika Scheynius, and Juha Kere. Differential DNA methylation in purified human blood cells: implications for cell lineage and studies on disease susceptibility. PloS one, 7(7):e41361, 2012.

[35] W Evan Johnson, Cheng Li, and Ariel Rabinovic. Adjusting batch effects in microarray expression data using empirical Bayes methods. Biostatistics, 8(1):118–127., 2007.

[36] Jovana Maksimovic, Lavinia Gordon, and Alicia Oshlack. Swan: Subset-quantile within array normalization for illumina infinium humanmethylation450 beadchips. Genome Biology, 13(6):R44, 2012.

